# Large scale discovery of coronavirus-host factor protein interaction motifs reveals SARS-CoV-2 specific mechanisms and vulnerabilities

**DOI:** 10.1101/2021.04.19.440086

**Authors:** Thomas Kruse, Caroline Benz, Dimitriya H. Garvanska, Richard Lindqvist, Filip Mihalic, Fabian Coscia, Ravi Teja Inturi, Ahmed Sayadi, Leandro Simonetti, Emma Nilsson, Muhammad Ali, Johanna Kliche, Ainhoa Moliner Morro, Andreas Mund, Eva Andersson, Gerald McInerney, Matthias Mann, Per Jemth, Norman E Davey, Anna K Överby, Jakob Nilsson, Ylva Ivarsson

**Affiliations:** The Novo Nordisk Foundation Center for Protein Research, University of Copenhagen, Faculty of Health and Medical Sciences, Blegdamsvej 3B, 2200 Copenhagen, Denmark; Department of Chemistry - BMC, Uppsala University, Box 576, Husargatan 3, 751 23 Uppsala, Sweden; Department of Clinical Microbiology, Umeå University, 90185 Umeå, Sweden; Laboratory for Molecular Infection Medicine Sweden (MIMS), Umeå University, 90186 Umeå; Department of Medical Biochemistry and Microbiology, Uppsala University, Box 582, Husargatan 3, 751 23 Uppsala, Sweden; Department of Microbiology, Tumor and Cell Biology, Karolinska Institutet, Stockholm, Sweden; Division of Cancer Biology, The Institute of Cancer Research, 237 Fulham Road, London SW3 6JB, UK

**Keywords:** viral hijacking, short linear motif, phage display, stress granule, G3BP, SARS-CoV-2

## Abstract

Viral proteins make extensive use of short peptide interaction motifs to hijack cellular host factors. However, current methods do not identify this important class of protein-protein interactions. Uncovering peptide mediated interactions provides both a molecular understanding of viral interactions with their host and the foundation for developing novel antiviral reagents. Here we describe a scalable viral peptide discovery approach covering 229 RNA viruses that provides high resolution information on direct virus-host interactions. We identify 269 peptide-based interactions for 18 coronaviruses including a specific interaction between the human G3BP1/2 proteins and an ΦxFG peptide motif in the SARS-CoV-2 nucleocapsid (N) protein. This interaction supports viral replication and through its ΦxFG motif N rewires the G3BP1/2 interactome to disrupt stress granules. A peptide-based inhibitor disrupting the G3BP1/2-N interaction blocks SARS-CoV-2 infection showing that our results can be directly translated into novel specific antiviral reagents.

---

RNA viruses such as the Ebola, dengue and coronaviruses cause a variety of diseases and constitute a continuous threat to public health. The coronaviruses are the largest single stranded RNA viruses known and their genomic RNA encodes around 30 viral proteins^1^. During infection each viral protein performs unique functions and interacts with a range of cellular protein host factors to allow viral proliferation and immune escape^2–5^. Precise disruption of viral-host factor interactions is an attractive strategy for developing novel antiviral reagents. The advantage of targeting these interactions is that resistance is less likely to develop and furthermore as the same host factor can be used by multiple viruses such reagents may provide broader spectrum activity. For this reason, numerous large scale mass spectrometry-based interaction screens^2, 3, 5^ as well as CRISPR based screening approaches^6–10^ have been used to uncover host factor interactions and dependencies for SARS-CoV-2 allowing repurposing of drugs^11, 12^. Although these methods have been transformative in our understanding of SARS-CoV-2 biology the molecular detail provided by these methods is not always sufficient to readily transform the results into novel antiviral reagents. Experimental approaches that would complement the existing powerful methods and provide a more detailed view of viral interactions with host factors could accelerate development of new antivirals.

An attractive class of protein interactions that can be inhibited for therapeutic purposes are viral short linear interaction motifs (SLiMs) that bind to defined pockets on globular domains of the host factor^13, 14^. SLiMs are short peptide motifs in unstructured regions of proteins and contain 2-3 amino acid binding determinants within a 10 amino acid stretch^15, 16^. Viruses extensively use SLiMs to hijack cellular host factors and SLiMs can readily evolve through mutations in unstructured regions allowing viruses to interact with novel host factors^17–19^. Despite the importance of SLiMs for understanding viral biology they are not uncovered by current methods^15, 16^. Proteomic peptide-phage display (ProP-PD) provides the opportunity to identify novel SLiM-based interactions and binding sites at amino acid resolution^20^. As shown in a small scale pilot study on C-terminal peptides of viral proteomes, it can be used to faithfully capture SLiM-based host-pathogen interactions^21^. Here we describe a novel phage-based viral peptide library to map SLiMs from 229 RNA viruses (Riboviria) mediating host factor interactions (Fig. 1A). This approach allows the simultaneous pan-viral identification of SLiM-based interactions with amino acid resolution of the binding sites. We document the power of this approach by identifying novel SARS-CoV-2 specific SLiM mediated host factor interactions and directly translate our screening results into novel mechanistic insights and pinpoint a potential target for antiviral intervention.

**Figure 1.**
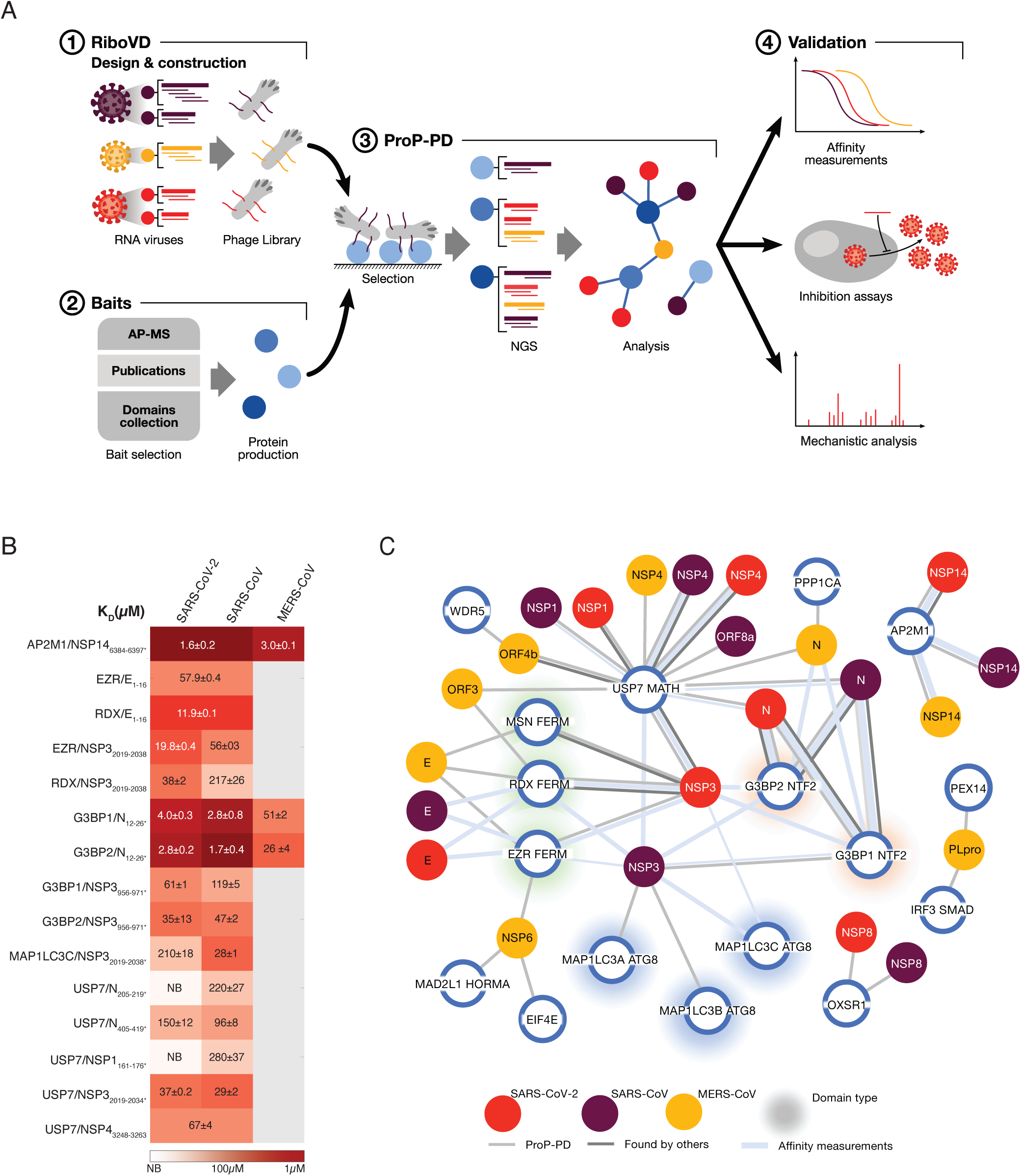
A pipeline for viral SLiM discovery. A) An overview of the platform for identifying viral SLiMs binding to cellular host factors. B) K_D_ values for the interactions between indicated viral peptides and host proteins. C) Network of SLiM mediated interactions between the indicated viral proteins from SARS-CoV-2 (red), SARS-CoV (brown) and MERS-CoV (yellow) and cellular host factors (blue circles). Light grey connecting line indicates interactions validated by affinity measurements, the weight of the line represents the affinity of the interaction (thick, 1-10 μM; medium, 11-100 μM; thin, 101-500 μM). Dark grey lines indicate protein-protein interactions with additional evidence found in the other studies (Table S2).

Exploiting recent developments in proteomic phage display (ProP-PD) technology^20, 22^ to identify RNA virus SLiMs binding to host factors. We designed a unique phage display library (RiboVD library) that tile the unstructured regions of 1074 viral proteins from 229 RNA viruses including SARS-CoV-2, SARS-CoV, MERS-CoV and 19 additional coronavirus strains. This library represents 19,549 unique 16 amino acid long peptides that are multivalently displayed on the major coat protein of the M13 phage. Given the recurrent pandemic potential of coronaviruses we used this library to understand interactions for this class of virus but note that it can be used for any RNA virus. We scanned published host factor interactomes for the SARS-CoV-2 viral proteins and recombinantly produced 57 domains from 53 cellular proteins reported to interact with SARS-CoV-2^2, 3, 5^. As transient SLiM based protein interactions might be lost during purifications of viral proteins for subsequent mass spectrometry analysis, we screened an additional set of 82 peptide binding domains, of which at least 27 have previously been reported to act as viral host factors and to be hijacked by SLiMs from viral proteins^23^. In total, 139 recombinantly expressed and purified human bait proteins (Table S1) were used in selections against the RiboVD library. Enriched phage pools were analyzed by next generation sequencing (NGS) to identify viral peptides that bound to the bait. This uncovered 269 putative SLiM-based interactions between 44 human protein domains (42 proteins) and 64 viral proteins from 18 coronavirus strains (Table S2). Of these, 117 (43%) interaction pairs involved human coronavirus proteins. We validated 27 out of 27 tested interactions using fluorescence polarization (FP) affinity measurements (Fig 1B, Fig S1, Table S3). We visualized the information generated for human and bat coronavirus proteins in an extensive network (Fig S2). We also generated a map of the viral proteins mediating SARS-CoV-2, SARS-CoV and MERS-CoV interactions with human host factors (Fig 1C). The map reveals common as well as unique interactions with host factors for these three coronaviruses. For instance, NSP14 of all three strains has a YxxL motif that binds to the clathrin coat adaptor protein AP2M1 with high affinity (Fig 1B; Fig S1, Table S3), which may be linked to trafficking of the viral protein or blocking of endocytosis of host proteins^24, 25^. The N-terminal region of the E protein from all three strains binds to the FERM domains of Ezrin and Radixin via a recently established [FY]x[FILV] SLiM^26^. Interestingly, our data show that the FERM domains also bind to NSP3 of SARS-CoV and SARS-CoV-2, thus, they can be targeted by distinct viral proteins. The SARS-CoV NSP3 FERM binding site overlaps with a [FWY]xx[ILV] binding site for the ATG8 domains of the autophagy related MAP1LC3A-C proteins. As an example of strain specific interactions, we found that an N-terminal peptide from the Nucleocapsid (N) proteins from SARS-CoV-2 and SARS-CoV bound to the NTF2 domain of the homologous G3BP1 and G3BP2 proteins (G3BPs) with high affinity (Fig 1B-C and S1, Table S3). This N peptide contains an ΦxFG SLiM (where Φ is a hydrophobic residue) that resembles motifs in USP10 and UBAP2L and in the alphavirus nsP3 protein known to bind a hydrophobic pocket in the NTF2 domain of G3BP^27–30^. The ΦxFG SLiM is also present in the N proteins from bat betacoronaviruses and consistently the corresponding bat HKU5 peptide was identified in our screen (Fig S2; Table S2).

To pinpoint therapeutically relevant host protein-viral SLiM interactions we screened three of the identified peptide motifs for antiviral activity. To this end, we generated lentiviral vectors expressing GFP fused to four copies of one viral SLiM reasoning that this would inhibit binding of the corresponding full-length SARS-CoV-2 protein to the specific host factors through competition. As a control we used GFP fused to SLiMs containing mutations in the binding motif. The host proteins targeted by viral peptides were G3BPs (SARS-CoV-2 N), Ezrin and Radixin (SARS-CoV-2 E and NSP3), and the MAP1LC3s (NSP3). VeroE6 cells were first transduced with the lentiviruses and 3 days later infected with SARS-CoV-2 and viral titer determined after 16 hours. This revealed that the G3BP-binding peptide from the N protein decreased viral titer 3.4-fold (Fig 2A). To obtain a more potent inhibition of the SARS-CoV-2 N-G3BP interaction, we used a 25 amino acid residue peptide from Semliki Forest vius (SFV) nsP3 containing two continuous FGDF like SLiMs that has previously been shown to bind G3PBs with high affinity^31^. Remarkably, this peptide binds approximately 10-fold stronger than the SARS-CoV-2 N peptide to both G3BP1 and G3BP2 (Kd= 4 μM vs Kd= 0.3 μM, Fig 2B and S3A). We constructed “G3BP inhibitors (G3BPi)” by fusing sequences encoding one or three copies of wild type (wt) or mutated (ctrl) SFV nsP3 SLiMs to GFP. As expected, mass spectrometry analysis confirmed that the major cellular targets of the G3BPi are the G3BPs (Fig 2C, S3B and Table S4). Furthermore, expression of the G3BPi wt but not G3BPi ctrl prevented binding of SARS-CoV-2 N to G3BP1 in cells (Fig 2D). Consistent with these binding and competition data, lentiviral mediated expression of the G3BPi in VeroE6 cells potently inhibited SARS-CoV-2 proliferation after 16 hours of infection (Fig 2A). An effect of the G3BPi was also evident in assays monitoring viral infection rates or replication (Fig 2E-F). In a cell based transfection assay monitoring assembly and release of virus-like particles mutating the G3BP binding motif in N had no effect (Fig S3D).

**Figure 2.**
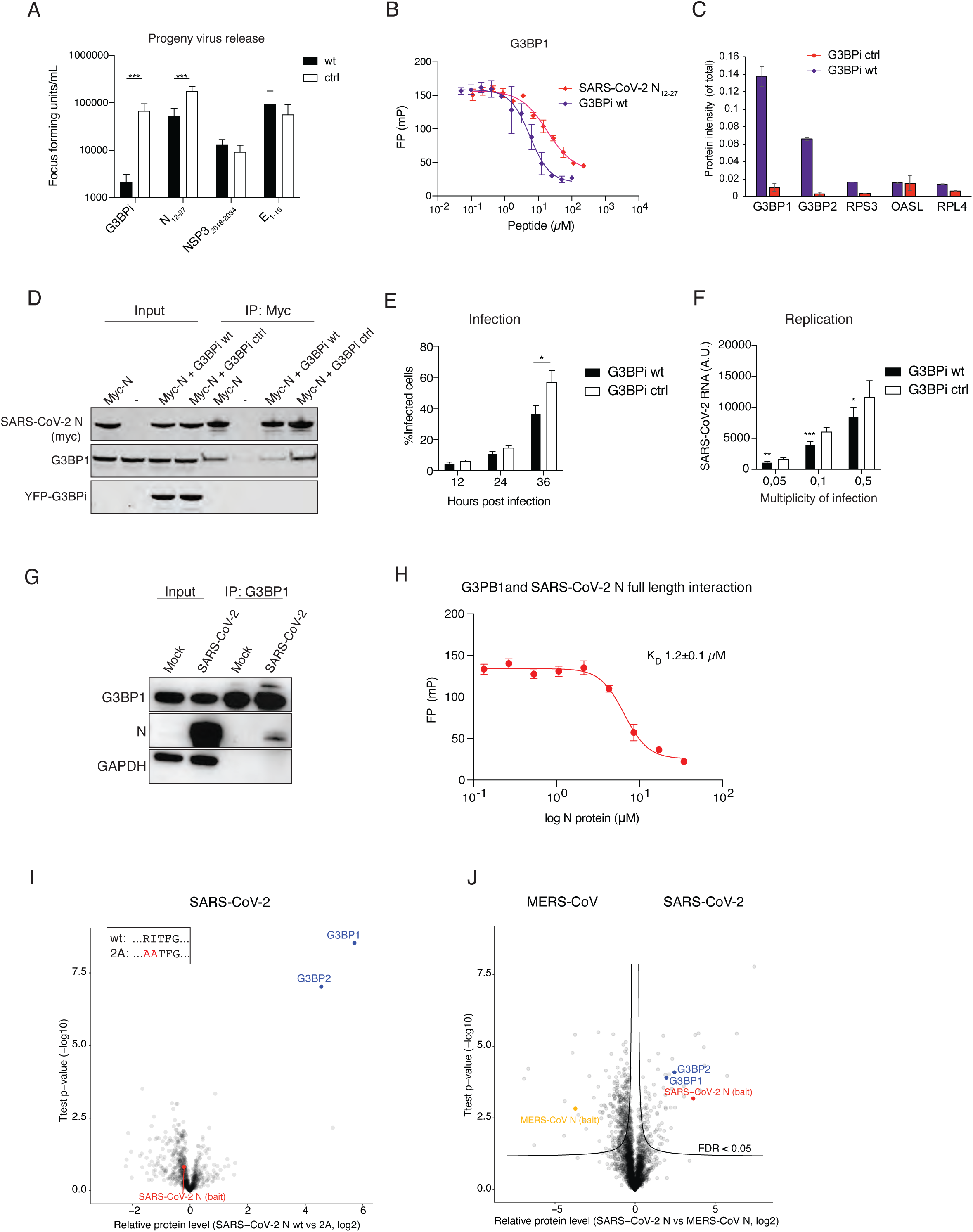
The interaction between N and G3BP1/2 is important for SARS-CoV-2 infection. A) Screen for SARS-CoV-2 antiviral activity of selected viral peptides as well as the G3BP inhibitor (G3BPi). The amount of SARS-CoV-2 virus released was determined 16 hours post infection by focus forming assay. B) Affinity measurements of recombinant G3BP1 NTF2 binding to G3BPi and the SARS-CoV-2 N peptide. C) Quantitative mass spectrometry comparison of G3BPi wt and ctrl purified from HeLa cells. D) Purification of myc-tagged SARS-CoV-2 N expressed in HeLa cells and its interaction with G3BP1 analyzed by western blot. G3BPi wt or ctrl were co-expressed with myc-tagged N where indicated. E) Effect of G3BPi on % infected VeroE6 cells during 36 hours of infection. F) Amount of SARS-CoV-2 RNA measured 16 hours post infection with qPCR at different MOI in VeroE6 cells expressing G3BPi or control inhibitor. G) Endogenous G3BP1 was purified from mock or SARS-CoV-2 infected cells and probed for N. H) In vitro interaction of recombinant full-length SARS-CoV-2 N and the NTF2 domain of G3BP1 as measured by fluorescence polarization spectroscopy. I) Quantitative mass spectrometry analysis of YFP tagged SARS-CoV-2 N wt or 2A purified from HeLa cells. J) Quantitative mass spectrometry analysis of YFP tagged N SARS-CoV-2 and N MERS purified from HeLa cells. Asterisks indicate statistical significance calculated by unpaired *T* test * *p*< 0.05, ** *p*< 0.01 and *** *p*<0.005

Thus, the approach presented here is useful for identifying important virus-host factor interactions which inhibit viral proliferation when disrupted.

The above results prompted us to further investigate the N-G3BP interaction and its function during infection. The coronavirus N protein is important for viral replication as well as packaging of the viral RNA^32–34^. The G3BPs are multi-functional RNA-binding proteins best known for their essential roles in innate immune signaling and the assembly and dynamics of cytosolic stress granules^35–38^. Stress granules are large protein-RNA assemblies formed in response to various stresses and viral infections^39–41^. Perhaps unsurprisingly, the G3BPs have turned out to be major targets for viral interference and several viral proteins have been shown to recruit G3BP1 to support viral replication and/or to inhibit stress granules formation^42^. Of note, the herpesviruses and alphaviruses have been shown to recruit G3BPs by SLiMs having resemblance to the sequence in N^28, 43–45^. However, a deeper mechanistic understanding for how viral proteins affect G3BP biology is missing. Given that the N-G3BP interaction was important for SARS-CoV-2 infection and presents a novel antiviral strategy we investigated this interaction in more detail. We first confirmed that the interaction between N and G3BP1 takes place in SARS-CoV-2 infected cells (Fig 2G). We also confirmed binding of recombinant full length N protein to G3BP1 using FP, which revealed an affinity similar to the N peptide (Fig 2H). To confirm the SLiM mediated interaction in cells, we compared the interactome of N wild type (N wt) to an N protein where we mutated two amino acids in the ΦxFG motif (N 2A) using label free quantitative mass spectrometry. This confirmed a highly specific N-G3BP1/2 interaction fully dependent on an intact ΦxFG motif (Fig. 2I and S3C, Table S4). Using a similar approach, we quantitatively compared the interactomes of the N protein from MERS-CoV with that of N from SARS-CoV-2, revealing specific binding of G3BPs to N from SARS-CoV-2 (Fig 2J, Table S4). This is in line with our observation that the N peptides containing the ΦxFG motif from SARS-CoV-2 and SARS-CoV bind to G3BPs with high affinity, while the corresponding MERS-CoV peptide bound weakly (Kd= 2.8 µM vs Kd= 26 µM, Fig 1B). Similar results were obtained by western blot, which also showed that the N protein from HKU1-CoV did not bind G3BPs, consistent with the ProP-PD results (Fig. S2 and S3E). Consistently, the ΦxFG motif resulted in the specific co-localization of mCherry tagged N protein from SARS-CoV and SARS-CoV-2 to arsenite induced stress granules in cells expressing YFP tagged G3BP1 (Fig 3A).

**Figure 3.**
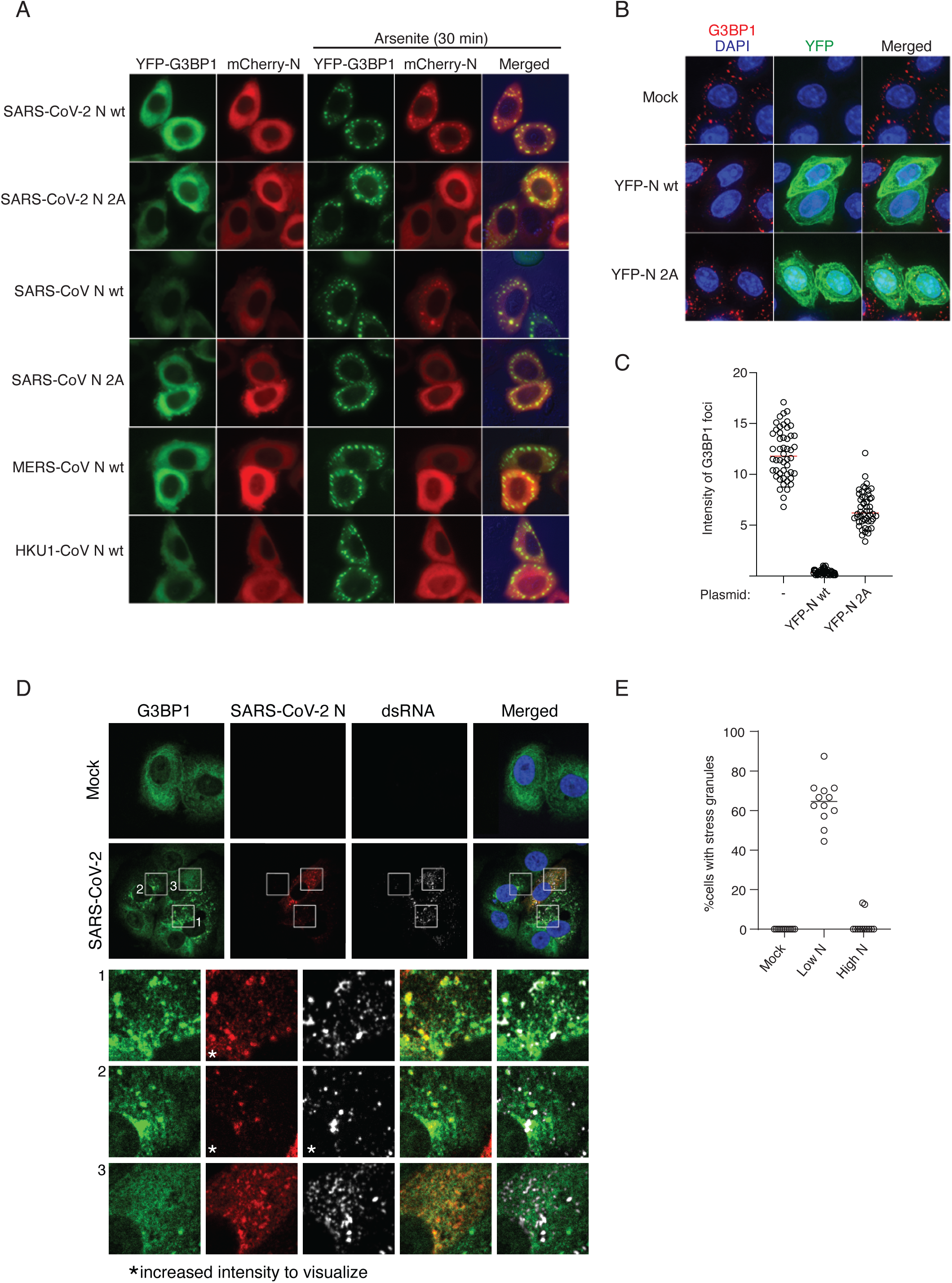
Interaction between SARS-CoV-2 N and G3BP1/2 affects stress granule formation. A) Live cell microscopy analysis of HeLa cells co-transfected with YFP-G3BP1 and mCherry tagged N proteins from the viral strains indicated. B) Effect of SARS-CoV-2 N wt and N 2A on arsenite induced stress granule formation as measured by immunofluorescence of endogenous G3BP1. C) Quantification of G3BP1 foci intensity from B). Red bar indicates median intensity, and each circle represents the intensity of one G3BP1 foci. At least five foci from 10 cells were measured. D) Immunofluorescence analysis of G3BP1, N and viral dsRNA in SARS-CoV-2 infected VeroE6 cells 6 hours post infection. E) Effect of N levels on G3BP1 foci formation in SARS-CoV-2 infected cells. Each circle represents the % of cells with high or low N levels that also express stress granules.

Recent publications have reported that SARS-CoV-2 N induces stress granule disassembly^46, 47^. but the mechanistic basis of this is unclear. To investigate the effect of the SARS-CoV-2 N ΦxFG motif on endogenous stress granule formation we overexpressed YFP tagged N wt or N 2A in HeLa cells and stained for endogenous G3BP1 following arsenite treatment. Quantifying the intensity of cytoplasmic G3BP1 foci in cells positive for YFP revealed that N WT expression disrupted stress granule formation more efficiently when the G3BP binding motif was intact (Fig 3B-C). Thus, the N-terminal ΦxFG motif of the N protein constitutes the main determinant of G3BP binding and stress granule disassembly. We next analyzed G3BP1 foci formation and cellular localization of viral dsRNA in relation to N protein expression levels in VeroE6 cells after six hours of SARS-CoV-2 infection (Fig 3D-E). At this timepoint, a mixture of early and later stage infected cells is observed. In mock treated cells we detected no cells with G3BP1 foci while infected cells with low levels (early-stage infection) of N protein had G3BP1 foci in about 60% of the cells (Fig 3E). In cells with low levels of N, this protein and viral dsRNA co-localized with G3BP1 to stress granules (Fig 3D). However, in cells with high levels of N protein (late-stage infection) we detected no G3BP1 foci. Collectively our results suggest that during early stages of infections the levels of N protein are insufficient to disrupt stress granule formation and instead N and viral dsRNA co-localize with G3BP1 in these structures. Once N concentrations reaches a certain threshold, this disrupts stress granules, and this depends on the ΦxFG motif. A possible interpretation of these observations is that during the earlier stages of infection SARS-CoV-2 takes advantage of the stress granule RNA machinery. Consistently, dsRNA and N co-localize with G3BP1 foci and when the N-G3BP interaction is inhibited a reduction of viral replication is observed (Fig 2F and 3D).

To understand how N could affect stress granule formation and G3BP function through the ΦxFG motif we set out to identify cellular G3BP interactors with similar binding motifs. To this end, we screened a novel ProP-PD library that displays the intrinsically disordered regions of the human proteome^22^ against the NTF2 domains from G3BP1 and G3BP2. The combined data set includes 72 peptides from 57 proteins with the majority of sequences containing a ΦxFG motif (Φ: [FILV]), thus resembling the sequence in the N protein (Fig 4A-B, Table S5). Nineteen of the proteins uncovered by the screen are in core stress granule proteins - including known peptide motifs in USP10 and UBAP2L, but also peptides from stress granule proteins that have not previously been reported to contain ΦxFG motifs. The screen also uncovered a peptide from Caprin-1 which has been shown to bind G3BPs but does not match the consensus sequence^48^ (Fig. 4A-B). This suggests that G3BPs serve as major hubs for stress granule biology in part by interacting with ΦxFG like motifs residing in several stress granule components. However, the screen also returned many peptides in proteins with roles outside of stress granule biology such as TRIM25 and IRF7 (anti-viral interferon signaling)^42^, and DDIT3 (endoplasmic reticulum stress)^49^. FP affinity measurements were used to confirm binding between the purified NTF2 domain of G3BP2 and several identified peptides originating from TRIM25, DDIT3, UBAP2L, Caprin-1, USP10 and PRRC2B (Fig S4A). Furthermore, we biochemically validated a number of the G3BP binding motifs in the context of the full-length proteins (Fig S4B). Caprin-1 and UBAP2L co-localized with G3BP1 in stress granules after arsenite treatment in a manner dependent on intact SLiMs (Fig S4C). Conversely, no stress granule localization was observed for TRIM25 and DDIT3 (Fig S4C) further supporting the notion that the G3BPs also have cellular roles beyond stress granule biology^50^.

**Figure 4.**
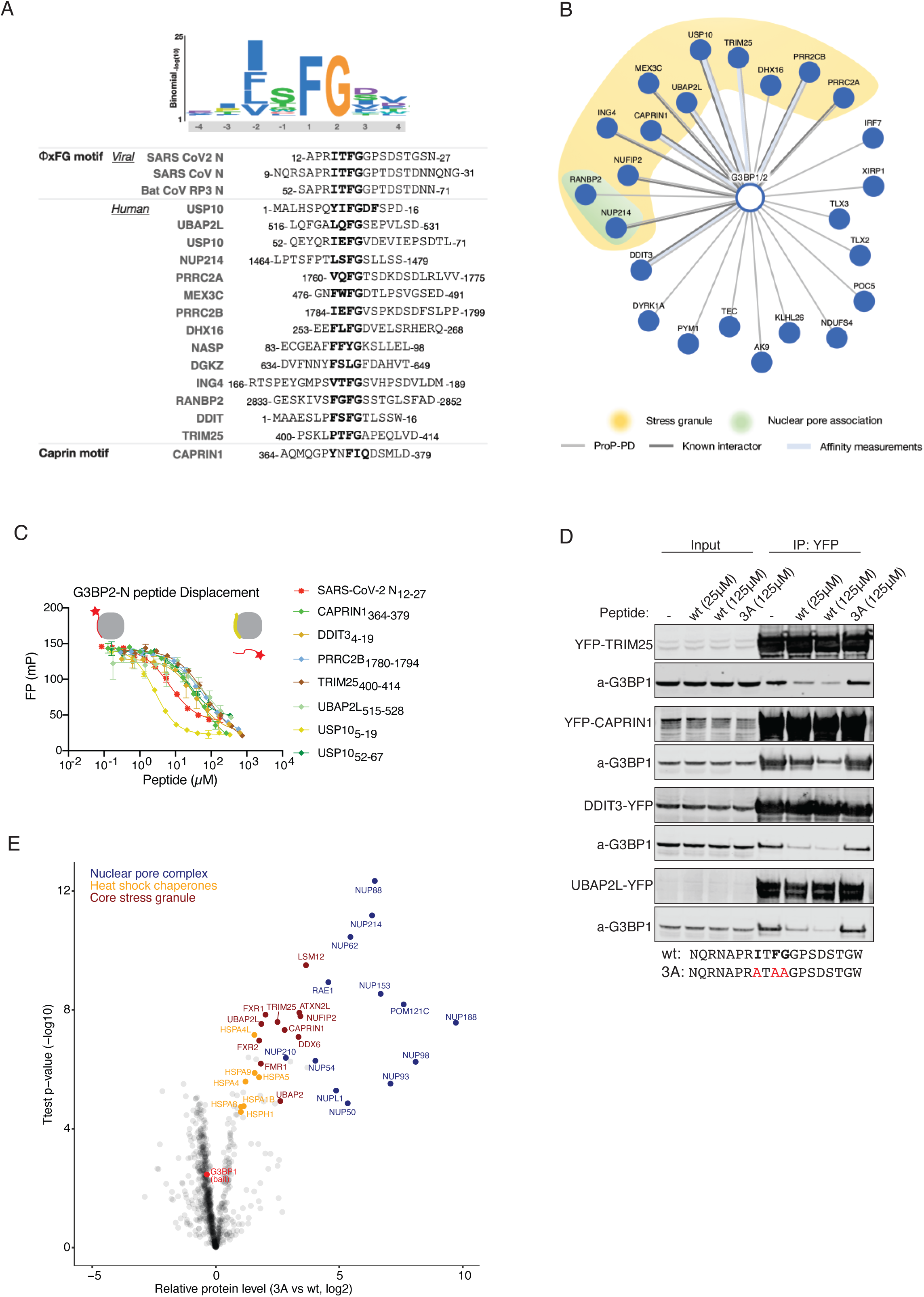
N competes with cellular proteins for binding to G3BP1 through its ΦxFG motif. A) Schematic of the position specific scoring matris ΦxFG for G3BPs NTF2 domains and sequence alignment of coronavirus peptides and selected human G3BPs ligands found through ProP-PD^17^. B) Network of a select set of human SLiM based interactions of the G3BPs found through ProP-PD. Light grey line indicates interactions validated by affinity measurements (Table S3), the weight of the line represents the affinity of the interaction (thick, 1-10 μM; medium, 11-100 μM; thin, 101-500 μM). Dark grey lines indicate protein-protein interactions with additional reported evidence. C) The SARS-CoV-2 ΦxFG N peptide competes with the indicated peptides for G3BP2 binding in vitro. D) The N ΦxFG peptide but not a control peptide competes with TRIM25, CAPRIN1, DDIT2 and UBAP2L for binding to G3BP1 in cells. E) G3BP1-YFP was affinity purified from HeLa cells in the presence of either a SARS-CoV-2 ΦxFG N wt or N 3A peptide and analysed by quantitative mass spectrometry.

The N protein is a highly expressed viral protein^51^ during infection so we hypothesized that it would compete with host cell proteins containing ΦxFG SLiMs for binding to G3BPs. Consistently, FP affinity measurements revealed competition between the N ΦxFG peptide and all of the 7 peptides we tested for interaction with G3BP2 (Fig 4C). Next, immunopurifications of full length YFP-tagged TRIM25, DDIT3, Caprin-1 and UBAP2L in the presence of either a N wt peptide or a N 3A peptide where the ΦxFG motif is mutated to AxAA were performed. As expected, the N wt peptide disrupted interactions to G3BP1 thus validating a direct competition between the N ΦxFG peptide motif and four G3BP binding proteins (Fig 4D). The observed competition between the viral N ΦxFG peptide and UBAP2L for binding to G3BP1 is particularly interesting since UBAP2L is required for stress granule assembly through a direct interaction to the G3BPs via its ΦxFG like motif^27, 29^. This suggests a mechanistic basis for the ability of the N protein to inhibit stress granule formation.

Given the high levels of N we speculated that it could mediate a general rewiring of the G3BP interactome through its ΦxFG motif. To test this on a global scale, we purified G3BP1-YFP from HeLa cells and added either N wt or the N 3A mutated peptide as competitors for cellular proteins. Quantitative label free mass spectrometry allowed us to determine the proteins being specifically displaced by the N wt peptide (Fig 4E and Table S4). This revealed specific displacement of 59 proteins including several core stress granule components. In addition, the N peptide also displaced a large number of nuclear pore complex components, heat shock chaperones of the Hsp70 family and proteins of the ASC-1 and CTLH complexes. Except the CTLH components all of these proteins have been reported to localize to stress granules^27, 52, 53^.

The displacement of nucleoporins from G3BP1 by the N peptide suggests that FG motifs, which are abundant in nucleoporins^54^ might recruit them to stress granules through direct interaction to the NTF2 domain of the G3BPs. Consistently, two ΦxFG like motifs from nucleoporins were selected in the G3BP ProP-PD screen (Figure 4A-B; Table S5). Importantly, Nup62, which we identify in our MS competition screen (Fig 4E) has been shown to be required for efficient SARS-CoV-2 infection^3^. It is possible that N displaces Nup62 from G3BPs to make it accessible for other viral processes. Together, we show that the N protein modulates the G3BP1/2 host interactome through its ΦxFG motif by competing with numerous cellular ΦxFG containing proteins. Our G3BP motif and mass spectrometric screens provide a rich resource for the future dissection of basic stress granule biology and G3BP signaling in general.

Collectively we describe a therapeutic relevant interaction between the ΦxFG SLiM in SARS-CoV-2 N and the G3BP proteins. Our results reveal that the N protein during infection hijacks G3BPs to viral replication centers likely to facilitate replication and possibly other aspects of viral RNA metabolism. The disruption of stress granules at later stages of infection could also dampen the cellular anti-viral response. Consistent with this idea we identify ΦxFG motifs in TRIM25, MEX3C and IRF7 that are key components of the G3BP-RIG-1 antiviral interferon pathway^42, 55–57^.

By screening the intrinsically disordered regions of 229 RNA viruses against a host factor in one go our ProP-PD screening approach uncovers both common principles shared by several viruses as well as interactions specific for a given strain. We show that the SLiMs can be screened for antiviral activity to pinpoint therapeutically relevant interactions. Given the amino acid resolution provided by the ProP-PD this can guide the development of agents targeting these interactions specifically. Our approach is scalable and easily applied to other relevant host factors and the library readily updated to incorporate novel RNA viruses emerging in the future. We thus foresee that this approach can be a powerful tool for future investigations of virus interactions with cellular host factors and for developing novel antivirals.

## Materials and Methods

### Design and construction of the RiboVD phage display library

The RiboVD library was designed using a previously described design pipeline^22^. The search space was defined as the UniProt reference proteomes of the mammalian and avian RNA viruses (Riboviria; taxonomic identifier: 2559587) and a representative proteome from RNA virus clades without a reference proteome (A complete list of the viral strains in the library is available at http://slim.icr.ac.uk/phage_libraries/rna_viruses/species.html). The accessible regions of the viral proteins were tiled as 16 amino acid long peptides overlapping by 12 amino acids as described^22^. The resulting library contained 19,549 unique peptides from 1,074 proteins (i.e. the numbers are based uncleaved polyprotein) in 229 viral strains across 211 families. The library design contains 4,172 coronavirus peptides mapping to 192 proteins from 23 strains including SARS-CoV-2, SARS-CoV and MERS-CoV-2. The complete library design is available at http://slim.icr.ac.uk/phage_libraries/rna_viruses/proteins.html.

The designed oligonucleotide library was obtained from a commercial provider (Genscript) and used to construct the RiboVD library that display the encoded peptides on the major coat protein P8 of the M13 phage following a published protocol^26^. In brief, the oligonucleotide library was used as template for 15 cycles amplification cycles of PCR amplification (98°C for 10 sec, 56°C for 15 sec and 72°C for 10 sec) using Phusion polymerase (Thermo Scientific) and primers complementary to the constant annealing regions flanking the designed library sequences. The PCR product was purified using a using a nucleotide removal kit (Qiagen), phosphorylated using T4 polynucleotide kinase (Thermo Scientific) for 1 h at 37°C and annealed to phagemid ssDNA (90°C for 3 min, 50°C for 3 min and 20°C for 5 min). dsDNA was synthesized using T7 DNA polymerase (Thermo Scientific) and T4 DNA ligase (Thermo Scientific) at 20°C for 16 h. The dsDNA was purified from a 1% agarose gel and electroporated into *E.coli* SS320 (Lucigen) electrocompetent cells pre-infected with M13KO7 helper-phage (ThermoFisher). The phages were allowed to propagate for 24 h in 2YT (10 g yeast extract, 16 g tryptone, 5 g NaCl per L) medium. The phage was precipitated from the supernatant by the addition of 1/5^th^ volume of 20% PEG8000/2.5 M NaCl and centrifugation at 27,000 x g for 20 min. The phage pellets were dissolved in phosphate-buffered saline (PBS, 137 mM NaCl, 2.7 mM KCl, 95 mM Na_2_HPO_4_, 15 mM KH_2_PO_4_ pH 7.5). The resulting phage library was re-amplified and stored at −80°C in the presence of 10% glycerol.

### Protein purification

*E.coli* BL21-Gold(DE3) (Agilent Technology) transformed with the plasmids encoding the His-GST-tagged proteins (Table S1) were grown in 500 mL 2YT at 37°C until an OD_600_ between 0.6 and 0.8. Protein expression was induced with 1 mM isopropyl ß-D-1-thiogalactopyranoside (IPTG) and allowed to proceed for 20 h at 18°C. The bacteria were harvested by centrifugation (4,500 g, 10 min). Proteins were batch purified using Glutathione Sepharose™ 4 Fast Flow Media (Cytiva,) or Ni^2+^ IMAC (Ni Sepharose™ 6 Fast Flow, Cytiva) following the manufacturer’s protocols. The purity of the proteins was validated via SDS-PAGE.

Proteins used for FP affinity measurements were expressed and purified in larger scale (at least 8 x 500 mL 2YT). His-GST tagged G3BP1 and G3BP2 were used directly for affinity measurements after dialysis. For the proteins, the tag was removed by cleaving with HRV 3C protease or thrombin, and further purified by reverse IMAC or by using a HiTrap Benzamindine FF (HS) (Cytiva) before being dialysed to a suitable buffer for FP affinity measurements (50 mM phosphate buffer, pH 7.5 and 1-2 mM reducing agent (DTT or TCEP)), except for RDX and EZR (20 mM HEPES pH 7.3, 150 mM NaCl, 3 mM DTT).

### Proteomic peptide phage selection (ProP-PD) and NGS analyis

Phage display selections against the immobilized bait proteins were performed in triplicate selections for four rounds of selection following a protocol described in detail elsewhere^26^. The RiboVD library constructed in this study was used in selections against 139 human bait proteins purified (Table S1). The NTF2 domains of G3BP1 and G3BP2 were further used in selections against a phage library that encodes the intrinsically disordered regions of the human proteome^22^. The peptide-coding regions of the naive phage library prior to any selection and the binding enriched phage pools (5 µL) were PCR amplified and barcoded using Phusion High-Fidelity polymerase (Thermo Scientific) for 22 cycles, using a dual barcoding strategy^58^ PCR products were confirmed through 2% agarose gel electrophoresis stained with GelRed, using a 50 bp marker (BioRad). The amount of the PCR products were normalized using Mag-bind Total Pure NGS (Omega Bio-tek) before pooling the samples. The resulting amplicon pool was further purified from a 2% agarose gel (QIAquick Gel extraction Kit Qiagen) with GelRed staining and eluted in TE (10 mM Tris-HCl, 1 mM EDTA. pH 7.5). The sample was analyzed using Illumina MiSeq v3 run, 1×150bp read setup, 20% PhiX by the NGS-NGI SciLifeLab facility. Results were processed using in-house custom Python scripts described previsouly^22^. Reads with an average quality score of 20 or more were kept, and their adapter regions and barcodes were determined allowing a maximum of 1 mismatch per adapter and/or barcode. The unambiguously identified reads were demultiplexed and their adapter and barcode regions were trimmed, Reads from each selection experiment were then translated into amino acid sequences and the number of counts for each unique peptide was determined.

To assess the quality of RiboVD library we analyzed the coverage percentage of the phage library over the library design using the NGS results of non-challenged naive library phage aliquotes. 95.50% of the peptide sequences designed to be in the library were confirmed by the NGS analysis of the naïve phage library.

### Analysis of the ProP-PD selection data

Following a recently outlined ProP-PD data analysis approach^22^ four metrics were used to rank the peptides i) NGS read counts, ii) peptide occurrence in replicated selections, iii) number of overlapped peptides, iv) motif match. The four metrics were then combined into a single score called ‘Confidence level’ forming 3 categories: high (4 metric criteria matched); medium (2 to 3 criteria matched); and low (only 1 metric is matched). Due to the relatively small size of the RiboVD library, the coronavirus dataset was further filtered for target specific ligands (occurring in less than 10 unrelated selections) (Table S2). The G3BPs HD2 P8 data was combined and a joint confidence score was calculated (Table S5). The peptide data was combined with available information on SG localized proteins from mass spectroscopy of purified mammalian stress granules^27, 52, 59^ and from other studies based on the information listed in the HIPPIE database. Each ProP-PD selected peptide was annotated with the above data and a count of the number of sources of evidence of SG localisation.

### PPI network visualization

Network’s visualization was done using Cytoscape^60^ using the data provided in Table S2 and Table S5.

### FP affinity determinations

FP experiments were carried out using a SpectraMax iD5 multi-mode microplate reader (Molecular Devices) using a black half area 96-well plate (Corning, USA #3993) with a total volume of 50 µL. The settings were set to 485 nm for excitation and 535 nm for emission. Peptides were from GeneCust (France) with a purity of over 95%. All measurements were performed at least in triplicates. For saturation experiment, proteins were serially diluted in the plate with phosphate buffer in a volume 25 µL, and then supplemented with a master mix of 10 nM FITC-labeled peptide in phosphate buffer. In the displacement assay, unlabeled peptides were serially diluted in the plate in 25 µL phosphate buffer, and 25 µL of displacement master mix was added (10 nM FITC-labeled peptide mixed with the protein of interest at a concentration 4 x the K_D_ determined through direct binding using the FITC-labeled probe peptide).

### Cell culture, virus and reagents

HeLa and HEK293 cells were maintained in DMEM GlutaMAX containing 100 U/ml penicillin, 100 mg/ml streptomycin, and 10% FCS (all from Thermo Fisher Scientific). Stable HEK cell lines were generated using the T-Rex doxycycline inducible Flp-In system (Invitrogen) and cultivated like HeLa cells with the addition of 5 μg/mL Blasticidin and 100 μg/mL Hygromycin B. VeroE6 cells were cultured in DMEM (Sigma) supplemented with 5% fetal bovine serum (FBS), 100 U/mL of penicillin and 100 μg/mL streptomycin (Gibco). The patient isolate SARS-CoV-2/01/human/2020/SWE accession no/GeneBank no MT093571.1, was provided by the Public Health Agency of Sweden. SARS-CoV-2 passage number 4 was cultured and titrated in VeroE6 cells. *E. coli* DH5*α* were maintained and propagated using standard microbiological procedures. The following drug concentrations were used: Sodium arsenite 0.5 mM, doxycycline 10 ng/ml unless otherwise stated.

### Expression constructs and cell line generation

Standard cloning techniques were used throughout. All N proteins and variants were generated by gene syntesis (Geneart). To generate the YFP-G3BPi inhibitor, the double FGDF motif from Semliki Forest virus nsP3 (RTTFRNKLPFTFGDFDEHEVDALASGITFGDFDDVL) or a control inhibitor (RTTFRNKLPATAGDFDEHEVDALASGITAGDADDVL) were fused to the C terminus of YFP by cloning the DNA encoding this into the pcDNA5/FRT/TO vector (Invitrogen). DNA encoding the G3BPi sequences was purchased from GeneArt, Life Technologies. All constructs were fully sequenced. The following point mutations were introduced using quick change mutagenesis to uncouple binding to G3BP: TRIM25 (F406A G407A), DDIT3 (F10A G11A), CAPRIN-1 (Y370S N371K F372S I373T), UBAP2L (F518L F523G). Detailed mutagenesis and cloning strategies are available upon request.

### Lentivirus production, transductions and virus infection

Lenti-X 293T cells (Takara bio) grown in a 100 mm plate format were co-transfected with 4.5 μg psPAX2 (Didier Trono lab, Addgene plasmid #12260), 500 ng pMD2.G (Didier Trono lab, Addgene plasmid #12259) and 5 μg of transfer plasmid per plate using polyethylenimine (Merck) as transfection reagent. At 24 h post transfection, medium was replaced with DMEM GlutaMAX containing 100 U/ml penicillin, 100 mg/ml streptomycin, and 10% FCS. Viral supernatant was harvested at 72 h post transfection, filtered through a 0.22-μm low protein-binding syringe filter and frozen at **−** 80°C. Transfer plasmids for lentiviral transduction were ordered from GenScript. To generate transfer plasmids, four copies of inhibitory peptide (three copies for the Semilikiforest peptide) or control peptides with the binding motifs mutated were fused to C terminus of EGFP and cloned to pLJM1-EGFP vector (David Sabatini lab, Addgene plasmid #19319).

For transductions VeroE6 or HEK cells were seeded into greiner CELLSTAR® 96-well plates or 6-well plates (VWR) containing lentivirus in DMEM containing 2 % FBS and 1 μg/mL polybrene, and incubated for 72 h. Transduced VeroE6 cells were infected with SARS-CoV-2 using the indicated MOI. Virus was detected at indicated time points using the same method as for viral titration except for using donkey anti-rabbit IgG Alexa Fluor 555 secondary antibody (Invitrogen). Nuclei were counterstained by DAPI. Number of infected cells were determined using a TROPHOS Plate RUNNER HD®. Number infected cells were normalized to DAPI count and presented as percentage infection of control peptide.

### Viral titration

Virus was diluted in ten-fold dilutions and added to VeroE6 cells followed by 1 h incubation at 37°C and 5% CO2. After 24 h of infection inoculum was removed and cells were fixed in 4 % formaldehyde for 30 minutes, permeabilized in PBS 0.5 % trition-X-100 and 20 mM glycine. Virus was detected using primary monoclonal rabbit antibodies directed against SARS-CoV-2 nucleocapsid (Sino Biological Inc., 40143-R001), and secondary anti-rabbit HRP conjugated antibodies (1:2000, Thermo Fisher Scientific). Viral foci were then stained by incubation with TrueBlue peroxidase substrate for 30 minutes (KPL, Gaithersburg, MD).

### Antibodies

The following antibodies were used at the indicated dilutions: c-Myc (1:1000, Santa Cruz Biotechnology, sc-40), rabbit anti-G3BP1 (WB; 1:1000, Cell Signaling Technology, #17798S), mouse anti-G3BP1 (IF; 1:1000, Abcam, ab56574), GFP-Booster_Atto488 (IF 1:300, ChromoTek), mouse anti-GFP (WB, 1:1000, Roche, 11814460001), rabbit anti-GFP (WB; 1:5000, in-house), rabbit anti-SARS-CoV-2 nucleocapsid (WB 1:2500; IF 1:500; Sino Biological Inc., 40143-R001), mouse APC-conjugated antibody directed against dsRNA J2 (IF 1:200, Scicons, 10010500) mouse anti-3xFlag M2 (WB 1:25000, Sigma, F1804), rabbit anti-Tubulin (WB 1:4000, Abcam, ab6046), donkey anti-rabbit IgG (Invitrogen), goat anti-rabbit HRP conjugated antibody (WB 1:2000 or 1:5000, Thermo Fisher Scientific, 31460), goat anti-mouse HRP conjugated antibody (WB 1:5000, Thermo Fisher Scientific, 31430), goat anti-mouse Alexa Fluor 546 (IF 1:1000, Invitrogen, A-11003), donkey anti-rabbit Alexa555 (IF 1:500, Thermo Fisher Scientific, a31572), donkey anti-mouse Alexa488 (IF 1:500, Thermo Fisher Scientific, a21202), goat IRDye 800CW anti-Mouse IgG (WB 1:10000, Li-Cor Biosciences, 926-32210), goat IRDye 800CW anti-Rabbit IgG (WB 1:10000, Li-Cor Biosciences, 926-32211), goat IRDye 680RD anti-Mouse IgG (WB 1:10000, Li-Cor Biosciences, 926-68070), goat IRDye 680RD anti-Rabbit IgG (WB 1:10000, Li-Cor Biosciences, 926-68071).

### RNA isolation, cDNA synthesis and qPCR

Total RNA (400 ng) was used to synthesize cDNA using High Capacity cDNA Reverse Transcription Kit (Applied Biosystems) according to the manufacturer’s instructions. GAPDH transcripts were detected by RT2 qPCR Primer Assay (Qiagen, Cat# 330001 PPQ00249A) and the qPCRBIO SyGreen Mix Hi-ROX kit (PCRBIOSYSTEMS), SARS-CoV-2 transcripts were detected using forward (GTCATGTGTGGCGGTTCACT) and reverse (CAACACTATTAGCATAAGCAGTTGT) primers and probe (FAM-CAGGTGGAACCTCATCAGGAGATGC-BHQ) and the qPCRBIO Probe Mix Hi-ROX kit (PCRBIOSYSTEMS). QPCR was run using a StepOnePlus fast real-time PCR system (Applied Biosystems).

### Live cell imaging

Live-cell analysis was performed on a Deltavision Elite system using a × 40 oil objective with a numerical aperture of 1.35 (GE Healthcare). The DeltaVision Elite microscope was equipped with a CoolSNAP HQ2 camera (Photometrics). Cells were seeded in eight-well Ibidi dishes (Ibidi) and before filming, the media was changed to Leibovitz’s L-15 (Life Technologies). Appropriate channels were recorded for the times indicated. For transient transfections, DNA constructs were transfected into HeLa cells using lipofectamine 2000 (Life Technologies) 24 h prior to analysis.

### Immunoprecipitations

Cells were lyzed in lysis buffer (150mM NaCl, 50mM Tris pH7.4, 1mM EDTA, 1mM DTT, 0,1% NP40) supplemented with protease and phosphatase inhibitors (Roche) for 25 min on ice. Lysates were cleared for 15 min at 20000 rcf and incubated with 20 μL pre-equalibrated GFP-trap or Myc-trap beads (ChromoTek) as indicated for 45 min at 4°C. Following 3 washes with lysis buffer, the beads were either eluted in 25 μL 2x LDS sample buffer (Novex, Life Technologies), boiled for 5 min, separated by SDS-PAGE and analyzed by Western Blotting with the indicated antibodies or subjected to quantitative mass spectrometry as described in the AP-MS section. For peptide competition experiments the indicated peptides were added to cell lysates for 30 min at 4°C before incubated with GFP-trap beads.

### Immunoprecipitation of SARS-CoV/SARS-CoV2/MERS-CoV N proteins

Two 15 cm^3^ dishes were seeded with HeLa cells at 20% confluency. On the following day, cells were transfected with 2.5 mg YFP-myc-N protein or YFP control plasmids. Cells were collected after 48h and lysed in 900mL lysis buffer: 100mM NaCl, 50mM Tris pH7.4, 0,1% IGEPAL (NP40), 1mM DTT supplemented with protease (Complete EDTA Free mini:Roche) and phosphatase (PhosStop: Roche) inhibitor tablets. Lysates were sonicated with Bioruptor for 10 cycles: 30s ON, 30s OFF intervals at 4°C and cleared at 20000 rcf for 45 min. The cleared lysate was incubated with pre-equilibrated GFP-Trap beads for 1h at 4°C and rotation. Three washes with 1mL wash buffer: 150mM NaCl, 50mM Tris pH7.4, 0,05% IGEPAL (NP40), 5% Glycerol, 1mM DTT, followed by one wash with 1mL basic wash buffer: 150mM NaCl, 50mM Tris pH7.4, 5% Glycerol. The supernatant was discarded, and beads were stored at −20°C before prepared for the MS analysis, or separated by SDS-PAGE followed by Western blot analysis.

### Immunofluorescence microscopy

For immunofluorescence microscopy, HeLa cell lines were seeded in eight-well Ibidi dishes (Ibidi) and transfected with the indicated constructs. 24 hours after transfection cells were treated with 0.5 mM sodium arsenite for 30 minutes to induce the formation of stress granules and subsequently fixed in 4% paraformaldehyde in PBS. Cells were blocked in 3% BSA in PBS-T for 30 min before incubation with GFP-Booster _Atto488 (1:300; Chromotek), or mouse anti-G3BP1 in 3% BSA in PBS-T for 1 h at room temperature. Unbound primary antibodies were removed by washing four times for 5 min in PBS-T at room temperature followed by incubation with secondary antibodies (Alexa Fluor 546; 1:1000; Invitrogen) and DAPI for 45 min. IBIDI dishes were then washed four times for 5 min in PBS-T. Z stacks 200 nm apart were recorded on a microscope (DeltaVision Elite) using a 40 × oil objective lens (numerical aperture 1.35) followed by deconvolution using SoftWoRx. Fluorescent intensity of stress granule signals was quantified by drawing a circle closely around stress granule signals and the intensity values from the peak continuous stacks were subtracted from the background of neighboring areas.

VeroE6 cells were seeded in 8-well chamber slides (Sarstedts) and infected with SARS CoV-2 for 6 h. Cells were fixed with 4% formaldehyde, quenched with 10 mM glycine, and permeabilized with PBS and 0.5% Triton X-100. Thereafter, cells are incubated with primary antibodies against SARS-CoV-2 nucleocapsid ((1:500) Sino Biological Inc., 40143-R001) and G3BP1 ((1:500) Abcam, ab56574) followed by incubation with conjugated secondary antibodies anti-rabbit Alexa555 and anti-mouse Alexa488 (1:500, Thermo Fisher Scientific). Then cells were stained with an APC-conjugated antibody directed against dsRNA J2 ((1:200) Scicons 10010500, antibody was conjugated using APC Conjugation Kit - Lightning-Link® (ab201807)). Nuclei were detected with DAPI (diluted 1:1500), coverslips were mounted and samples were analyzed using a Zeiss 710 (Carl Zeiss, Oberkochen, Germany) confocal microscope with a 63x oil objective (Zeiss) and ZEISS ZEN Imaging Software. Number of cells highly infected (strong N staining) was counted in 12 random fields (magnification 20x) and % of cells expressing stress granule (G3BP1) signals in these were calculated. The % of stress granulas in low N was calculated in a similar way, however, the N signal intencity was increased, to visualize the low levels of infection.

### Assembly assay

pcDNA3.1 expression plasmids coding for the structural proteins of SARS CoV-2, Spike (S D614), Membrane (M) (was a gift from Jeremy Luban (Addgene plasmid #158074 and 158078; http://n2t.net/addgene:158078; RRID: Addgene_158078)), Envelope 3xFlag (pGBW-m4252867 was a gift from Ginkgo Bioworks & Benjie Chen (Addgene plasmid # 153626; http://n2t.net/addgene:153626; RRID: Addgene_153626) and pcDNA5/FRT/TO Myc SARS CoV2 N and N 2A was transfected into HEK293T cells using GeneJuice (Novagen, Darmstadt, Germany) following the manufacturer’s protocol (12 μg of DNA/sample in total). Cells and Virus-Like Particles (VLPs) were collected 24 h after transfection. Cells were lysed (0.5 M Tris-HCl pH8, 1 M NaCl, 1% Triton X-100) and supernatant of transfected cells was collected, and concentrated by ultracentrifugation (100,000 × *g*, 90 min 4 °C, SW41, Beckman Coulter, Brea, CA). The pellet was resuspended in reducing Laemmli SDS-PAGE sample buffer. Proteins were separated with SDS-PAGE and Western blot analysis was performed using antibodies against SARS CoV-2 nucleocapsid, 3xFlag M2 (Sigma, F1804) and tubulin (Abcam, ab6046).

### Affinity purification and mass spectrometry (AP-MS)

Partial on-bead digestion was used for peptide elution from GFP-Trap Agarose (Chromotek). Briefly, 100 μl of elution buffer (2 M urea; 2 mM DTT; 20 μg/ml trypsin; and 50 mM Tris, pH 7.5) was added and incubated at 37°C for 30 min. Samples were alkylated with 25 mM CAA and digested overnight at room temperature before addition of 1% trifluoroacetic acid (TFA) to stop digestion. Peptides were desalted and purified with styrene–divinylbenzene reversed-phase sulfonate (SDB-RPS) StageTips. Briefly, two layers of SDB-RPS were prepared with 100 μl wash buffer (0.2% TFA in H2O). Peptides were loaded on top and centrifuged for 5 min at 500 g, and washed with 150 μl wash buffer. Finally, peptides were eluted with 50 μl elution buffer (80% ACN and 1% ammonia) and vacuum-dried. Dried peptides were dissolved in 2% acetonitrile (ACN) and 0.1% TFA in water and stored at −20°C.

### LC-MS analysis

Liquid chromatography mass spectrometry (LC-MS) analysis was performed with an EASY-nLC-1200 system (Thermo Fisher Scientific) connected to a trapped ion mobility spectrometry quadrupole time-of-flight mass spectrometer (timsTOF Pro, Bruker Daltonik GmbH, Germany) with a nano-electrospray ion source (Captive spray, Bruker Daltonik GmbH). Peptides were loaded on a 50 cm in-house packed HPLC-column (75 µm inner diameter packed with 1.9 µm ReproSilPur C18-AQ silica beads, Dr. Maisch GmbH, Germany). Peptides were separated using a linear gradient from 5-30% buffer B (0.1% formic acid, 80% ACN in LC-MS grade H2O) in 43 min followed by an increase to 60% buffer B for 7 min, then to 95% buffer B for 5min and back to 5% buffer B in the final 5min at 300nl/min. Buffer A consisted of 0.1% formic acid in LC-MS grade H2O. The total gradient length was 60 min. We used an in-house made column oven to keep the column temperature constant at 60 °C.

Mass spectrometric analysis was performed essentially as described in Brunner et al.^61^ in data-dependent (ddaPASEF) mode. For ddaPASEF, 1 MS1 survey TIMS-MS and 10 PASEF MS/MS scans were acquired per acquisition cycle. Ion accumulation and ramp time in the dual TIMS analyzer was set to 100 ms each and we analyzed the ion mobility range from 1/K0 = 1.6 Vs cm-2 to 0.6 Vs cm-2. Precursor ions for MS/MS analysis were isolated with a 2 Th window for m/z < 700 and 3 Th for m/z >700 in a total m/z range of 100-1.700 by synchronizing quadrupole switching events with the precursor elution profile from the TIMS device. The collision energy was lowered linearly as a function of increasing mobility starting from 59 eV at 1/K0 = 1.6 VS cm-2 to 20 eV at 1/K0 = 0.6 Vs cm-2. Singly charged precursor ions were excluded with a polygon filter (otof control, Bruker Daltonik GmbH). Precursors for MS/MS were picked at an intensity threshold of 1.000 arbitrary units (a.u.) and resequenced until reaching a ‘target value’ of 20.000 a.u taking into account a dynamic exclusion of 40 s elution.

### Data analysis of proteomic raw files

Mass spectrometric raw files acquired in ddaPASEF mode were analyzed with MaxQuant (version 1.6.7.0)^62, 63^. The Uniprot database (2019 release, UP000005640_9606) was searched with a peptide spectral match (PSM) and protein level FDR of 1%. A minimum of seven amino acids was required including N-terminal acetylation and methionine oxidation as variable modifications and cysteine carbamidomethylation as fixed modification. Enzyme specificity was set to trypsin with a maximum of two allowed missed cleavages. First and main search mass tolerance was set to 70 ppm and 20 ppm, respectively. Peptide identifications by MS/MS were transferred by matching four-dimensional isotope patterns between the runs (MBR) with a 0.7-min retention-time match window and a 0.05 1/K0 ion mobility window. Label-free quantification was performed with the MaxLFQ algorithm^64^ and a minimum ratio count of two.

### Bioinformatic analysis

Proteomics data analysis was performed with Perseus^65^ and within the R environment (https://www.r-project.org/). MaxQuant output tables were filtered for ‘Reverse’, ‘Only identified by site modification’, and ‘Potential contaminants’ before data analysis. Missing values were imputed after stringent data filtering and based on a normal distribution (width = 0.3; downshift = 1.8) prior to statistical testing. For pairwise proteomic comparisons (two-sided unpaired t-test), we applied a permutation-based FDR of 5% to correct for multiple hypothesis testing including an *s_0_* value^66^ of 0.1.

## Author contributions

ND designed the RiboVD library, MA constructed it. CB, MA, FM, EA and JK purified proteins. CB and YI performed phage selections. LS and ND analyzed NGS results. YI and AS analyzed PPI data. CB, FM, MA and JK performed affinity measurements and analyzed data. AMM analyzed G3BP:N interaction in virus infected cells. RTI generated lentiviral constructs and particles. RL performed viral infections and microscopy, inhibition assays, EN performed assembly assays and microscopy. TK performed analysis of stress granules in HeLa cells and analysis of G3BP motifs and competition studies. DHG prepared samples for N interactomes and western blot analysis of N purifications. JN performed cloning of N expression constructs. FC, AM, MM performed mass spectrometry analysis. ND, PJ, AÖ, JN and YI conceived the study. YI, JN, AÖ supervised the project. JN and TK drafted initial manuscript and all authors contributed to the final version.

## Data Availability

The mass spectrometry proteomics data have been deposited to the ProteomeXchange Consortium via the PRIDE partner repository^67^ with the dataset identifier PXD025410.

## Acknowledgments

This work was supported by the grants from the Swedish Foundation for Strategic research (YI, PJ: SB16-0039), the Swedish Research Council (YI: 2020-03380; PJ: 2020-04395; AÖ: 2017-02438, 2018-05851; GM: 2020-01541), the Knut and Alice Wallenberg Foundation (grants to Science for Life Laboratory, 2020.0182), and by a grant from Sygeforsikring Danmark (2020-0176 to JN, MM, YI, GM) and by Cancer Research UK (ND: C68484/A28159). We thank the medical faculty Umeå University strategic research resource and the Laboratory for Molecular Infection Medicine Sweden for generous support (AÖ), and the Biochemical Imaging Center at Umeå University and the National Microscopy Infrastructure, NMI (VR-RFI 2016-00968) for assistance in microscopy. Sequencing was performed by the SNP&SEQ Technology Platform in Stockholm. The facility is part of the National Genomic Infrastructure (NGI) Sweden and Science for Life Laboratory and is also supported by the Swedish Research Council and the Knut and Alice Wallenberg Foundation. Work at the Novo Nordisk Foundation Center for Protein Research is supported by grant NNF14CC0001.

We thank Prof. Sachdev Sidhu for providing constructs for expression of 59 proteins (PMID: 33438817), Dr. Andreas Ernst ATG8 domain constructs (PMID: 28028054), and Prof. Carlos Fontes and Renaud Vincentelli for constructs for PDZ domain expression (PMID:31267466). pGEX Grb2 SH2-SH3 was provided by Bruce Mayer (Addgene plasmid # 46442), GST-Hsp90 N(9-236) by William Sessa (Addgene plasmid # 22481), intersectin I SH3 A domain by Peter McPherson (Addgene plasmid # 47413), pGEX-5X-1-LARP7 by Blerta Xhemalce (Addgene plasmid # 113545), pGEX-2T PTP-1B by Ben Neel (Addgene plasmid # 8602), pGEX-4T-1-RIPK3 by Jaewhan Song (Addgene plasmid # 78827), pGEX6P-1-SF2 by Honglin Chen (Addgene plasmid # 99020), pGEX VAMP7 (1-188) by Thierry Galli (Addgene plasmid # 42315), the SARS-CoV-2 pcDNA 3.1 SARS-CoV-2S D614, and M by Jeremy Luban (Addgene plasmids #158074, #158078), pGBW-m4252867 expressing E 3xFLAG by Ginkgo Bioworks & Benjie Chen (Addgene plasmid #153626). We thank the protein production and characterization platform at NNF CPR for producing full length SARS-CoV-2 N protein. We thank Josephine K. Duel for help with preparation of samples for MS analysis.

---

**Figure S1.**
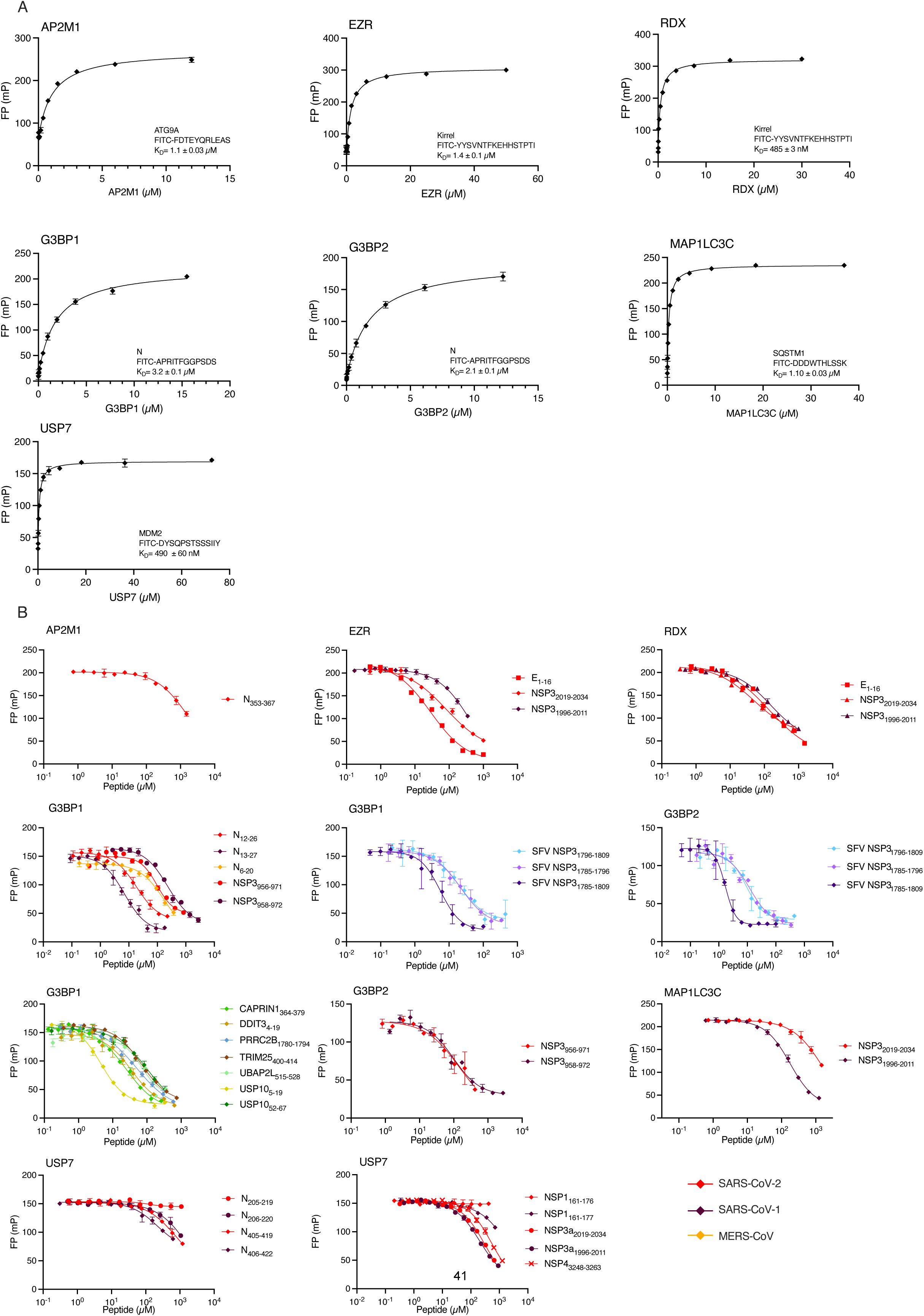
FP affinity determinations. A) Determination of K_D_ values of the FITC-labelled probe peptides. B) Affinity determinations through FP competition experiments using unlabeled peptides. All experiments were performed in triplicates.

**Figures S2.**
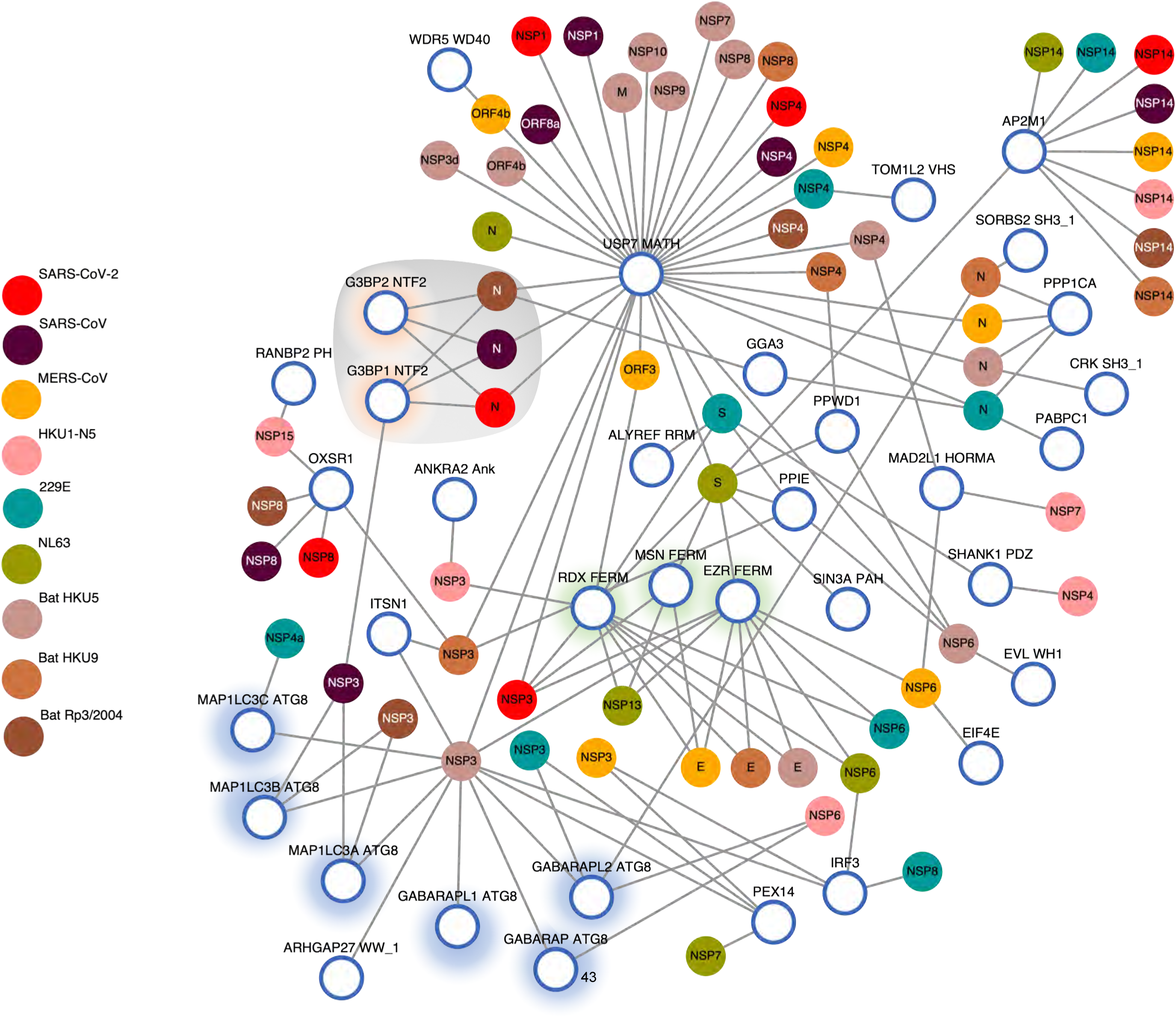
Network of coronavirus interactions with host factors. Interactions between indicated coronavirus (human and bat) proteins and host factors based on the results from the ProP-PD screen.

**Figure S3.**
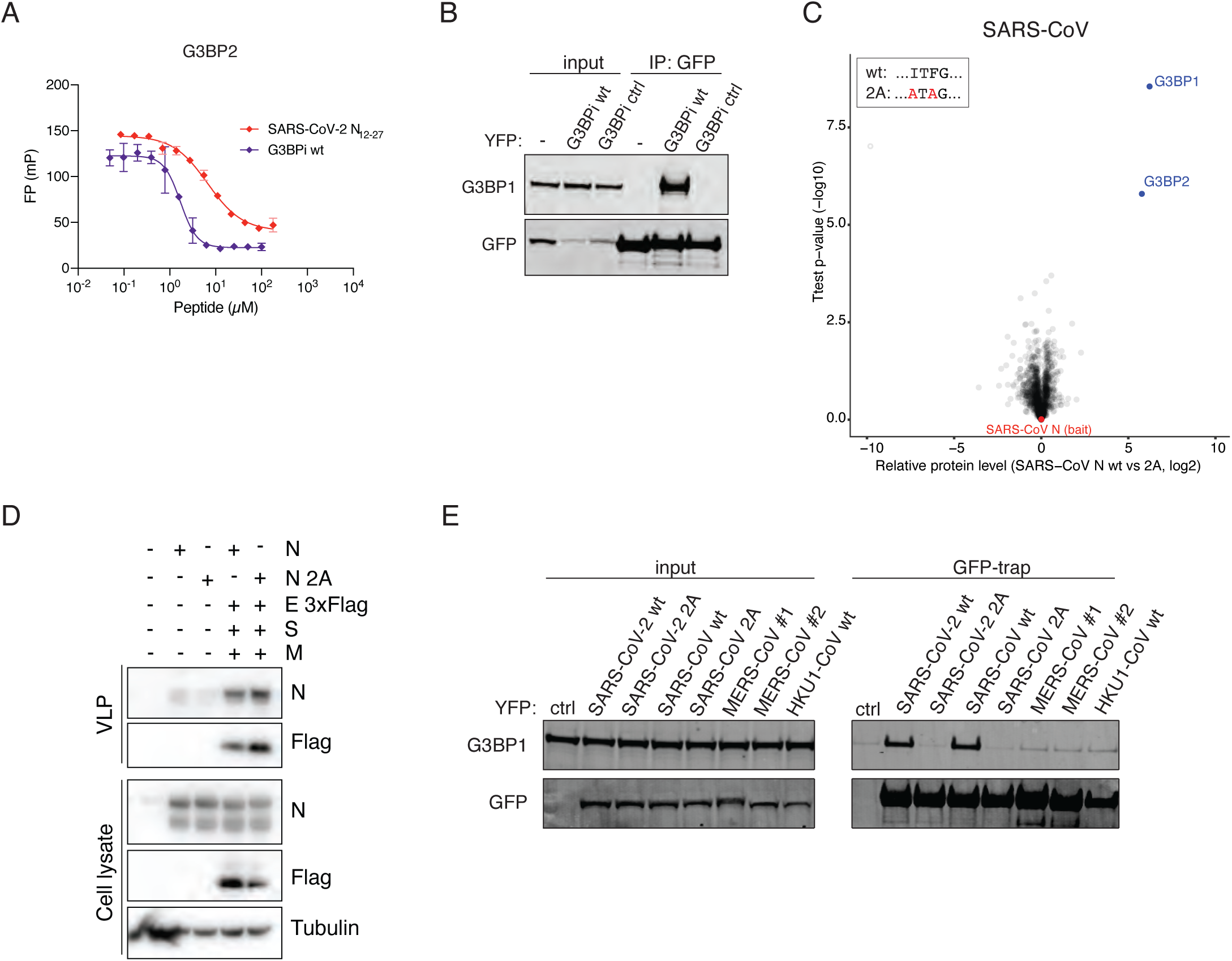
Analysis of G3BP-N interaction. A) In vitro affinity measurements between recombinant G3BP2 NTF2 and the SFV nsP3 and SARS-CoV-2 N peptides as indicated. B) Purification of YFP tagged G3BPi wt and ctrl from HeLa cells and probed for binding to G3BP1 by Western blotting. C) Quantitative mass spectrometry comparison of YFP tagged SARS-CoV N wt and 2A purified from HeLa cells. D) Virus assembly assay comparing the release of virus–like particles containing N wt and N 2A. E) YFP affinity purification of indicated N proteins from HeLa cells and probed for G3BP1 binding. Two different strains of MERS (#1 and #2) were analysed due to differences in sequence at the N terminal region of N.

**Figure S4.**
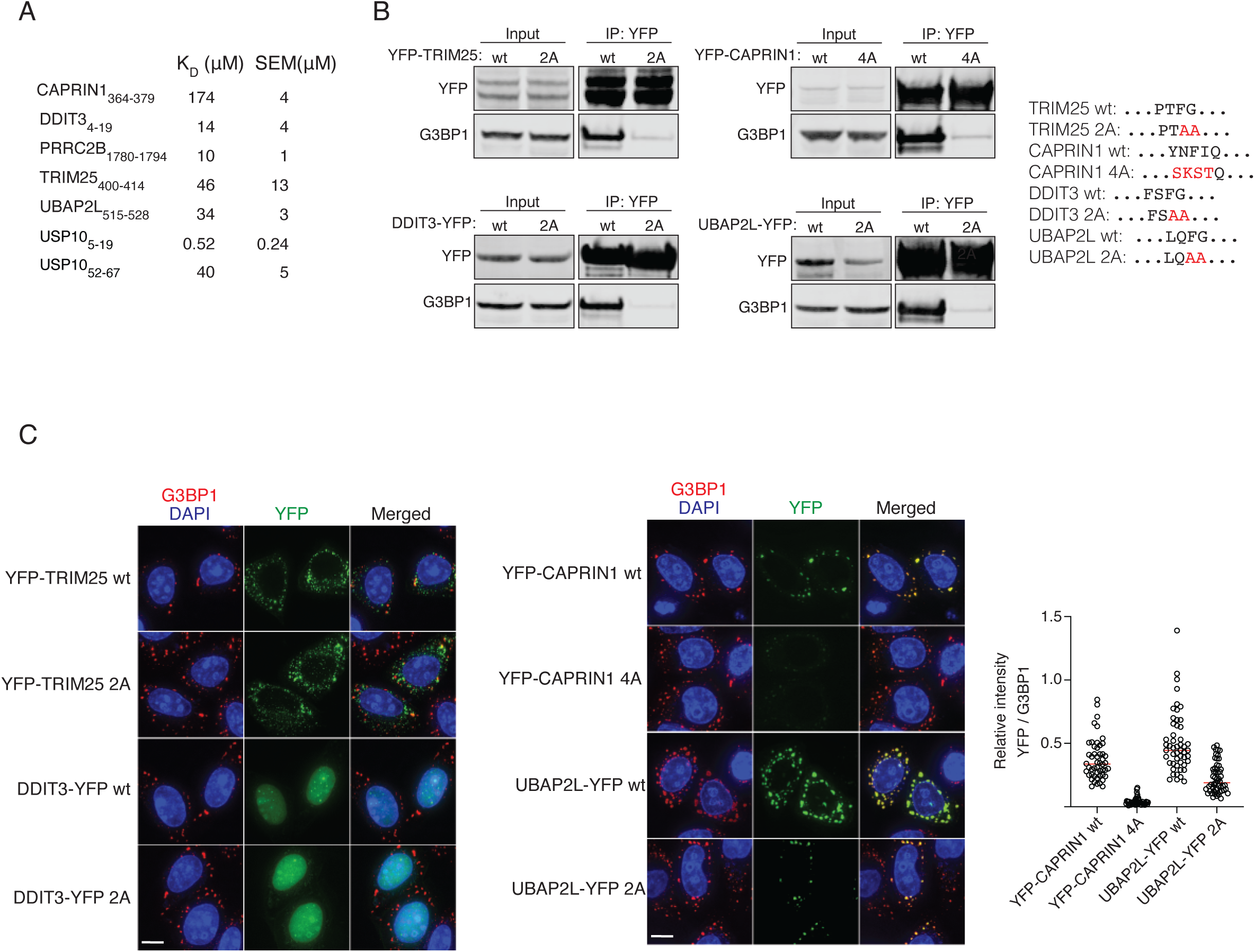
Validation of ΦxFG motifs in human proteins identified by ProP-PD. A) K_D_ measurements of indicated peptides binding to G3BP2. B) YFP affinity purification of indicated YFP tagged proteins either wild type (wt) or with mutated G3BP1 binding motif (2A or 4A) and probed for G3BP1 binding. C) Analysis of the indicated proteins and their co-localisation with G3BP1 following arsenite treatment for 30 minutes by immunofluorescence microscopy. The co-localization of CAPRIN1 and UBAP2L with G3BP1 into foci was quantified. Red bar indicates median relative YFP/G3BP1 intensity, and each circle represents one co-localization event. At least five foci from 10 cells were measured.

## Overview of supplemental tables

Table S1 Table of expression constructs and information on proteins screened

Table S2 Table of viral ProP-PD results

Table S3 Table of measured affinities including peptide information

Table S4 Mass spectrometry data

Table S5 G3BP1/2 ProP-PD selections

